# Continuous Seizure Emergency Evoked in Mice with Pharmacological, Electrographic, and Pathological Features Distinct from Status Epilepticus

**DOI:** 10.1101/2021.05.18.444686

**Authors:** Kevin M. Knox, Dannielle K. Zierath, H. Steve White, Melissa Barker-Haliski

## Abstract

**Objectives:** Benzodiazepines are the standard of care for the management of sustained seizure emergencies, including status epilepticus (SE) and seizure clusters. Seizure clusters are a variably defined seizure emergency wherein a patient has multiple seizures above a baseline rate, with intervening periods of recovery, distinguishing clusters from SE. While phenotypically distinct, the precise pathophysiological and mechanistic differences between SE and seizure clusters are under studied. Preclinical interrogation is needed to help uncover the behavioral, physiological, and pathological mechanisms associated with seizure emergencies in order to better manage these events in the susceptible individual.

**Methods:** Herein, we characterize a novel model of sustained seizure emergency induced in CF-1 mice through the combined administration of high-dose phenytoin (PHT; 50 mg/kg, i.p.) and pentylenetetrazol (PTZ; 100 mg/kg, s.c.).

**Results:** In the present manuscript we describe a mouse model of sustained seizure emergency that is physiologically, pharmacologically, and histologically distinct from SE. Acute administration of PHT 1 hour prior to s.c.PTZ led to significantly more mice with continuous seizure activity (CSA; 73.4%) versus vehicle-pretreated mice (13.8%; p<0.0001). CSA was sensitive to lorazepam and valproic acid when administered at seizure onset, as well as 30-minutes post-seizure onset. Carbamazepine worsened seizure control and post-CSA survival. Mice in CSA exhibited EEG patterns distinct from kainic acid-induced SE and s.c.PTZ alone, clearly differentiating CSA from SE and s.c.PTZ-induced myoclonic seizures. Neuropathological assessment by FluoroJade-C staining of brains collected 24-hours later revealed no neurodegeneration in any mice with CSA, whereas there was widespread neuronal death in brains from KA-SE mice.

**Significance:** This study defines a novel mouse model on which to elucidate the mechanistic differences between sustained seizure emergencies (i.e. SE and seizure clusters) to improve discovery of effective clinical interventions and define mechanisms of seizure termination.

**Key Points Box:**

- Seizure clusters are a variably defined seizure emergency that is sensitive to benzodiazepines, distinct from status epilepticus.
- The mechanistic differences between seizure clusters and status epilepticus are not well defined.
- We report a mouse seizure emergency model that is phenotypically, pathologically, and pharmacologically distinct from status epilepticus.
- This mouse model provides a novel platform on which to further interrogate the mechanisms underlying seizure emergencies.

## Introduction

Although the majority of seizures are brief, extensively long seizures require pharmacological intervention. Seizure emergencies include status epilepticus (SE) and seizure clusters. Clinically, SE is defined as a continuous seizure lasting >30 minutes or two or more sequential seizures without intervening recovery of consciousness.^1^ SE affects 50,000-150,000 people in the United States each year, with associated adult mortality ranging from 10-30%.^2; 3^ Between 4-16% of patients with epilepsy will have at least one SE event.^4^ Conversely, seizure clusters constitute a seizure emergency broadly defined by acute episodes wherein seizure control is interrupted by recurrent seizures lasting <5 minutes with periods of seizure freedom.^5; 6^ This seizure cessation and onset cycle uniquely differentiates clusters from SE. However, a lack of consensus clinical definition may lead to great variability in clinical prevalence.^7^□ Further, cluster seizures are a medical emergency unique to patients with epilepsy, whereas SE can occur in any individual, highlighting a key pathogenic difference.

The pharmacological sensitivity of each seizure emergency depends on a multitude of factors. Seizure clusters remain acutely sensitive to benzodiazepines (BDZs) even when drug administration is delayed.^8–11^ Conversely, SE can quickly become refractory to BDZs, leading to increased morbidity and mortality.^1; 12–14^ Yet, if left untreated, cluster seizures can rapidly progress to SE,^15–17^ highlighting conserved mechanistic similarities in two otherwise distinct indications.

Numerous models of SE exist that reproduce the clinical facets of SE, including behavioral and electrographic features, BDZ resistance, high seizure-induced mortality, and neurodegeneration.^18; 19^ Kainic acid (KA)-induced SE in rodents produces a sustained behavioral and electrographic seizure that is refractory to BDZs and causes neuronal death throughout the limbic system.^20^ Rodent models of KA-induced SE have been instrumental in understanding pathogenic mechanisms,^21^ the biological underpinnings,^20; 22^ and identifying novel interventions.^23^ Importantly, animal models of spontaneous recurrent seizures, including those that arise post-SE, do experience “cluster” seizures.^24; 25^ However, there is yet no predictable way to reliably produce cluster seizures in epilepsy models because of the high degree of interindividual variability.^26^ It has been thus challenging to consistently reproduce all types of clinical seizure emergencies in preclinical models to help elucidate the pathophysiological processes that differentiate SE from seizure clusters.

Acute administration of high-doses of sodium channel-blocking antiseizure drugs (ASDs), including phenytoin (PHT), can be proconvulsant and reduce seizure threshold in preclinical models.^27; 28^ Previous studies suggested that high-dose PHT pretreatment in the subcutaneous pentylenetetrazol (s.c.PTZ) model of myoclonic seizures in mice results in continuous seizures that were sensitive to midazolam and thus by *de facto* represented SE.^29^ Yet, the phenotypic characterization of this model has been notably absent. In combination with our serendipitous observations and prior publications demonstrating a proconvulsant effect of high-dose PHT,^27–29^ we sought to define whether the seizure induced by administration of high-dose PHT in the s.c.PTZ model would generate a sustained seizure with a distinguishable phenotype or pathology.^18; 27^ Mice pretreated with high-dose PHT prior to s.c.PTZ exhibit worsened seizure severity and duration; we thus hypothesized that this drug combination and resulting aberrant continuous seizure activity (CSA) represented a preclinical seizure model that is behaviorally similar to, yet pharmacologically distinct from, SE models. The present investigation thus sought to establish the phenotypic, electrographic, and neuropathologic features of this model of CSA. Like other SE models, the present CSA model is suitable for high-throughput drug screening to potentially identify new and improved treatments for non-SE seizure emergencies. Further, our present study provides an innovative preclinical platform on which to clarify the mechanistic basis of sustained seizures, seizure termination, and neurodegeneration after CSA, which may improve the management of seizure emergencies in the clinical setting.

## Methods

### Animals

Male CF-1 mice (25-40 g; 5-6 weeks old; Envigo) were housed 5 mice/cage with corncob bedding in a temperature-controlled specific pathogen-free vivarium on a 14:10 light/dark cycle (lights on: 6h00, lights off: 20h00). Animals were given access to irradiated chow (Picolab 5053) and filtered water, except during periods of behavioral manipulation. Mice were allowed to acclimate to the housing facility for at least 5 days and to the testing room for at least 1 hour prior to behavioral manipulation. All studies were conducted during the animals’ light phase. Animals were euthanized by CO_2_ asphyxiation. This study was not designed to assess the impact of sex as a biological variable, thus only male mice were used. All animal use was approved by the University of Washington Institutional Animal Care and Use Committee, conformed to the ARRIVE Guidelines,^30^ and was conducted in accordance with the United States Public Health Service’s Policy on Humane Care and Use of Laboratory Animals.

### Chemical Compounds

PTZ (catalog #P6500), PHT (catalog #PHR1139), methylcellulose (VEH; catalog #M0430), valproic acid (VPA; catalog #P4543), carbamazepine (CBZ; catalog #C4024), potassium permanganate (KMnO_4_; catalog #223468), xylenes (catalog #XX0060), xylazine (X1251-1G), and DPX mounting medium (catalog #06522) were all from Sigma Aldrich (St. Louis, MO, USA). Lorazepam (LZP; NDC 0641-6046-01) was from Westward Pharmaceuticals. Formal Fixx (catalog #9990244), Hoeschst 33342 (catalog #62249), and NaOH (catalog #SS254) were from Thermo Fisher Scientific (Waltham, MA, USA). FluoroJade-C (FJ-C; catalog #1FJC) was from Histo-Chem Inc (Jefferson, AR, USA). Kainic acid (KA; catalog #0222) was from Tocris (Bristol, UK). Ketamine (Ketaset) was from Zoetis, Inc (Parsippany, NJ).

### Induction of Continuous Seizure Activity

Each mouse was pre-treated with either 50 mg/kg PHT or VEH administered by the i.p. route (n=8/pre-treatment group). One hour later, 100 mg/kg PTZ was administered s.c. to each mouse which was then placed in an individual observation chamber (7.6H x 10.1W x 12.7L cm). At this dose, the seizure associated with s.c.PTZ is characterized by hindlimb extension and differs phenotypically from the minimal clonic seizure endpoint typically evaluated in drug intervention studies.^31^ The latency to onset of the PTZ-induced seizure, characterized by first twitch, clonic seizures, and onset of CSA was recorded for each mouse. Onset of CSA was visually defined as the time when the mouse jumped violently and then exhibited continuous forelimb and hindlimb clonus. Body weights were recorded 24 hours post-CSA and brains were collected for subsequent histology.

### Assessing the Impact of Pharmacological Intervention on Continuous Seizure Activity

To define whether CSA was phenotypically distinct from acute SE, we employed two ASD intervention paradigms (all i.p.): 1) immediate intervention (administration of ASDs within ≤1 minute CSA onset); or 2) delayed intervention (administration of ASDs 30 minutes post-CSA onset). Once a mouse entered CSA, it was randomized to receive VEH (0.5% methylcellulose), CBZ (20 or 40 mg/kg), LZP (2 or 4 mg/kg), or VPA (150 or 300 mg/kg). Each mouse was then monitored for 1 hour and all seizure activity was recorded. There were n=8 mice/group, which gives 80% power at 95% significance to detect a 1.5 standard deviation difference in seizure severity between each group versus VEH with p<0.05.s Low and high doses of VPA and CBZ were chosen as those that can block a maximal electroshock (MES) tonic hindlimb extension seizure in 50% of mice (ED50) and produce rotarod impairment in 50% of mice (TD50), respectively.^32; 33^ The dose of LZP was selected based on our ED50 in CF-1 mice in the MES test (1.20 mg/kg [95% CI: 0.801-1.79 mg/kg]; not shown) and was consistent with efficacy of other BDZs in the MES and rotarod tests.^32; 33^

### Kainic Acid-Induced Status Epilepticus

SE was induced with KA according to a modified method for C57BL/6 mice.^34^ Male CF-1 mice (n=10 for histology; n=3 for EEG) were treated with an initial 10 mg/kg (i.p.) dose of KA, followed by an additional 5 mg/kg (i.p.) dose of KA every 20 minutes until the presentation of multiple Racine stage 4 or 5 generalized seizures. Seizures were recorded during the SE induction period and for 1 hour after the onset of secondarily generalized behavioral seizures. Brains of mice that survived for 24 hours after onset of KA were collected by flash-freezing on dry ice.

### Surgical Implantation of EEG Electrodes for Video-EEG Monitoring

A separate cohort of mice (n=12) was used for video-EEG (vEEG) recordings. Mice were anesthetized with ketamine/xylazine (100/10 mg/kg; i.p.) and placed in a stereotaxic instrument.

A midline incision of approximately 1-2 cm was made exposing the skull. A 3-channel stainless steel electrode unit (MS333/1-A; Plastics One, Roanoke, VA) was placed in 2 holes drilled bilaterally, 4 mm posterior and lateral to bregma to overlay hippocampus. The third electrode was placed as a ground posterior to lambda to overlay cerebellum, a region known to be electrically silent. Three stainless steel mounting screws (00-96 × 1/16; InVivo1, Roanoke, VA) were placed, 2 posterior and lateral to the hippocampal electrodes and 1 anterior and lateral to bregma.^35^ The entire assembly was anchored with dental cement (Lang, 1223CLR). Mice were allowed to recover from anesthesia on a heating pad until ambulatory, and given oral Rimadyl (Bio-Serve, MD150-2) as an analgesic (0.5 mg). All implanted mice were individually housed and allowed to recover for 1 week after surgery.

### Video-EEG Monitoring of Electrographic Seizures

On the vEEG acquisition day, implanted mice (n=12) were connected to EEG electrodes and allowed 1 hour to acclimate to the recording room before obtaining a 1-hour baseline. Implanted mice were randomized into one of three experimental groups: VEH+PTZ (n=3; i.e., s.c.PTZ control group); KA-induced SE (n=3); or PHT+PTZ induced CSA (n=6). Recording of behavioral and electrographic seizures continued for >2 hours after onset for each experimental condition.

Video-EEG was recorded using a customized data acquisition system with an MP160 and EEC 100 (Biopac, Goleta, CA) and 3-channel electrodes (InVivo1).^26; 35^ The electrographic seizure analysis was conducted offline following completion of in-life studies using the baseline and active seizure period according to the following methods: the average power of two, 10 min EEG recording bins was first defined from the 1-hour baseline period. Then, the average power of two, 10 min active EEG recording bins was defined from the 1-hour period surrounding the onset of active VEH+PTZ induced clonic seizures, KA-induced SE, or PHT+PTZ induced CSA. The 10 min recording period for both baseline and active seizures was binned in 30 second epochs by Assyst EEG analysis software (KaosKey; Sydney, Australia). The time of onset of clonic seizures, SE, or CSA was visually noted by a trained investigator and the EEG spectrum corresponding to the 1-hour period immediately following the onset of behavioral seizures analyzed for each mouse.

### FluoroJade-C Assessment of Post-Insult Neurodegeneration

A separate cohort of mice (n=10 for KA-SE; n=16 for CSA) were euthanized 24 hours after seizure onset and brains removed, flash frozen, and maintained at −80°C until processing. Two consecutive 20 μm-thick sections/mouse from dorsal hippocampus (AP from Bregma: −1.34mm) and ventral hippocampus (AP from Bregma: −2.24mm) were sectioned on a cryostat (Leica DM1860) and slide-mounted. Slides were fixed in 4% Formal Fixx (10 minutes) and rinsed in DI water followed by FJ-C and Hoechst staining.^36; 37^ Slides were imaged on an upright fluorescent microscope (Leica DM-4) with a 10x objective (40x final magnification) with acquisition settings held constant. Images (n=4/mouse) were analyzed by an investigator blinded to treatment and visually scored for the presence of FJ-C positive cells in limbic structures (CA1, CA3, and dentate gyrus (DG) of dorsal hippocampus) and hypothalamic nucleus. If 2 or more of the images from each mouse demonstrated FJ-C positive staining in the designated brain region, the animal was considered to have FJ-C positive labeling.

### Statistics

The dose of PTZ necessary to elicit sustained twitch and clonus in >99% of mice (convulsant dose, CD99) was calculated by Probit.^38; 39^ Weights and latency to any seizure (in seconds) for acute pharmacological studies were analyzed by one-way ANOVA. The number of mice displaying CSA in acute pharmacological studies and mortality rates were compared by Fisher’s exact test. Survival after CSA onset was determined with a Kaplan-Meier plot. With the exception of CD99, all statistical analysis was performed in GraphPad Prism version 6.0 or later, with p<0.05 considered significant.

## Results

### Pentylenetetrazol and Phenytoin Dose Responses

We first sought to establish the dose of s.c.PTZ necessary to reliably induce seizures in >99% of mice (CD99), as well as the dose of PHT necessary to elicit CSA with low mortality when administered in combination of s.c.PTZ. The CD99 of s.c.PTZ was 102 mg/kg [95% CI: 83.4-220], thus 100 mg/kg PTZ was used for further studies (Figure 1B). We then quantified the dose-related impact of PHT on the conversion to CSA following administration of 100 mg/kg of PTZ (Figure 1D); pretreatment of mice with high dose PHT (50 mg/kg) induced CSA rather than typical myoclonic seizures, consistent with previous findings.^29^ Thus, subsequent behavioral, pharmacological, and electrographic studies were conducted with this combination; i.e., 50 mg/kg i.p. PHT administered 1 hour before 100 mg/kg s.c.PTZ.

**Figure 1.**
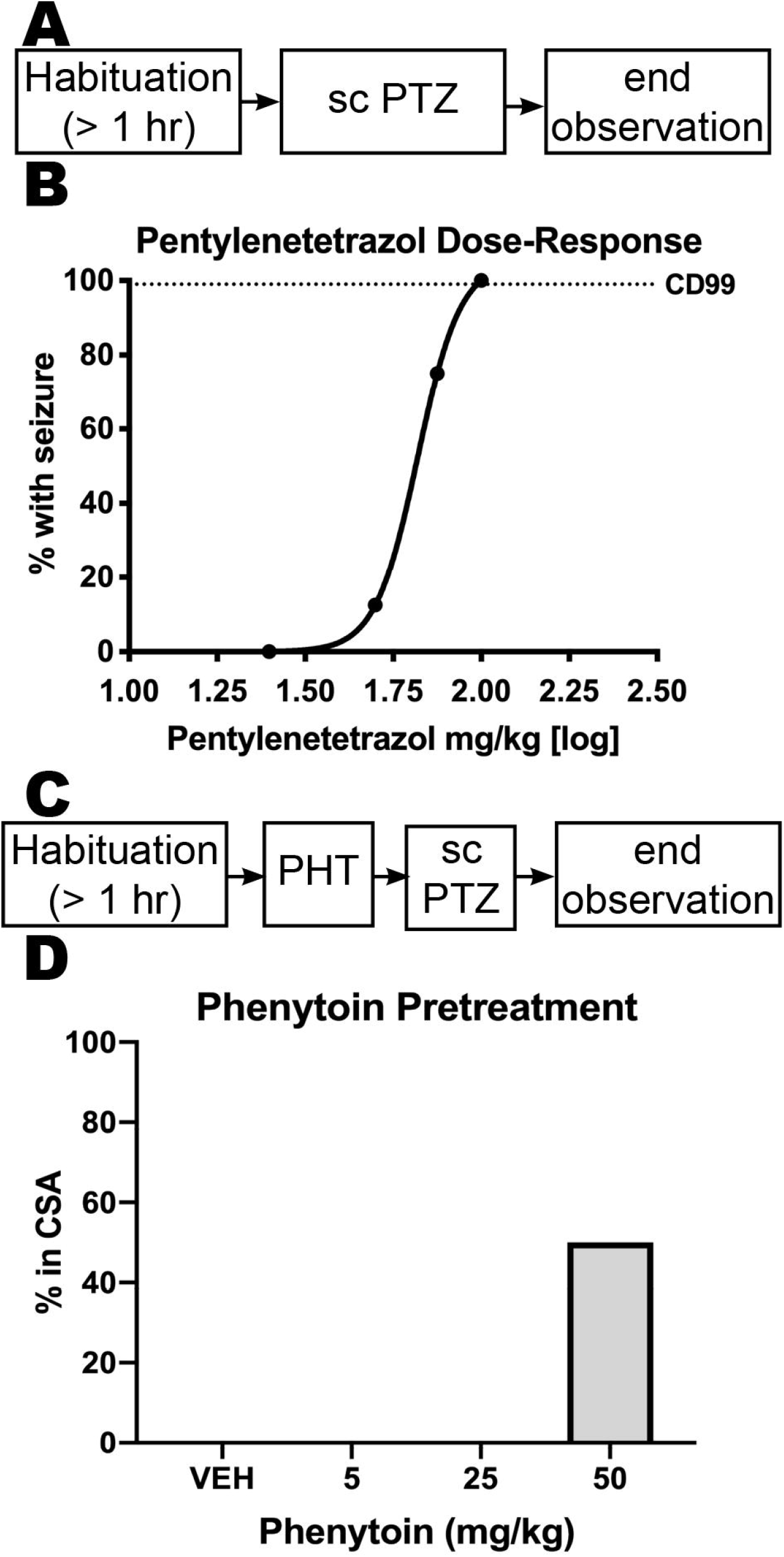
Pentylenetetrazol (PTZ) administered by the subcutaneous (s.c.) route normally induces clonic seizures in male CF-1 mice in a dose-related manner. This clonic seizure was shifted to continuous seizure activity (CSA) with pre-administration of high dose intraperitoneal (i.p.) phenytoin (PHT). (A) Timeline of dose-response studies to determine convulsant doses of PTZ in male CF-1 mice. Mice were allowed to habituate to the testing environment for 1 hour prior to administration of escalating doses of s.c.PTZ. B) The doses of s.c.PTZ were administered until at least two points could be established between the limits of 0% and 100% of mice with seizures to define the convulsant dose of 99% of mice (CD99). The CD99 of s.c.PTZ was determined to be 102 mg/kg. C) Timeline of the experimental design used to determine the pretreatment dose of i.p. PHT that would result in CSA in male CF-1 mice treated with s.c.PTZ (100 mg/kg). (D) Administration of 50 mg/kg PHT 1 hour prior to s.c.PTZ resulted in CSA in at least 50% of tested mice (n=4), which aligned with our pilot studies to develop this model (not shown) and previously published work.^29^ Administration of lower doses of PHT 1 hour prior to s.c.PTZ administration does not result in CSA onset in any mice.

### Phenytoin Pretreatment Precipitates Onset of Continuous Seizure Activity

Of the 94 mice pretreated with 0.5% MC one hour prior to 100 mg/kg s.c.PTZ, 13/94 (13.8%) developed seizures characteristic of CSA (Figure 2A). In contrast, 179/244 mice (74%) pretreated with high dose PHT (50 mg/kg) one hour prior to 100 mg/kg s.c.PTZ developed CSA (Fisher’s exact test, p<0.0001; Figure 2A). We then characterized the progression of CSA and assessed whether CSA significantly influenced mortality across time in the delayed intervention studies (Figure 2B). Of the 118 mice enrolled in the delayed intervention studies after PHT+PTZ induced-CSA, 64/118 (54.2%) survived to the 30-minute window to become candidates for pharmacological intervention. Of the mice that did not survive until the 30-minute intervention window, the median latency to mortality after CSA onset in the absence of pharmacological intervention was 10 minutes (Figure 2B). Thus, there was a notable selection bias for mice that survived the CSA insult to the 30-minute intervention point. If a mouse survived to the 30 min intervention window, it was generally more likely to survive, which may have also contributed to the discrepancy in improved survival between VEH-pretreated animals in the immediate and delayed intervention studies (Table 1).

**Figure 2.**
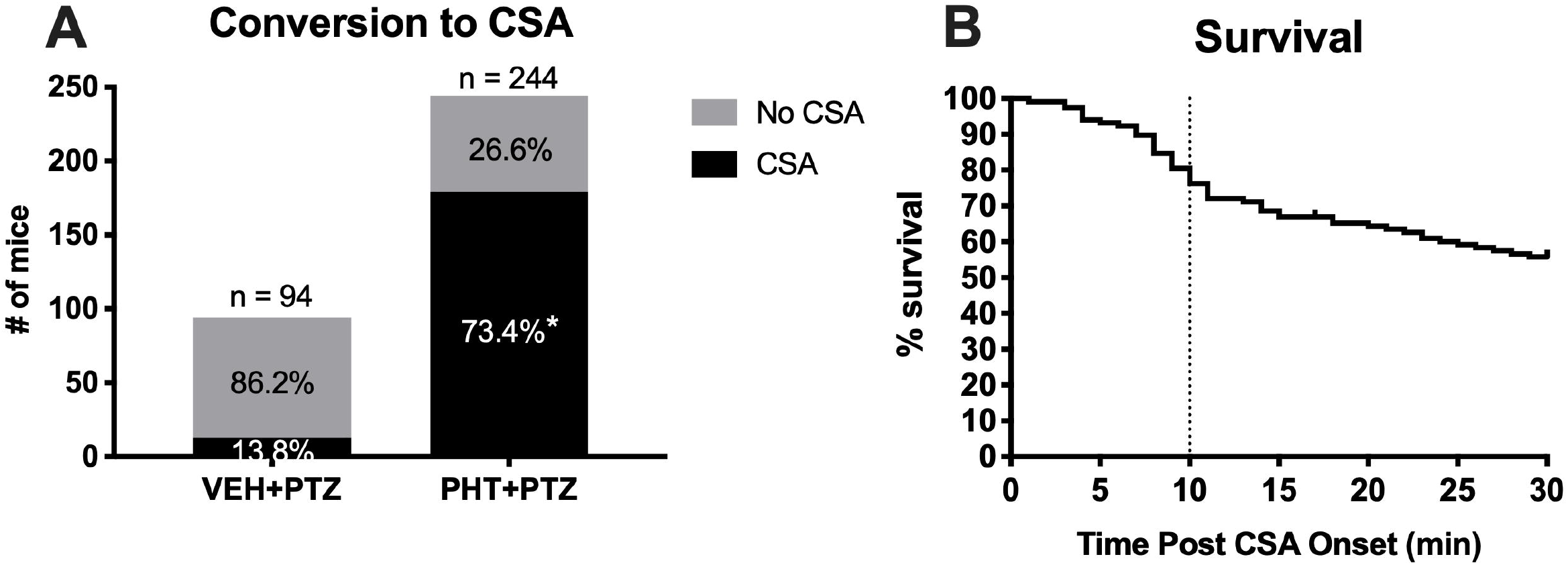
The combined administration of phenytoin (PHT; 50 mg/kg, i.p.) 1 hour prior to s.c. administration of pentylenetetrazol (PTZ) greatly increases the number of male CF-1 mice that present with continuous seizure activity (CSA). (A) There is a significant increase in the conversion to CSA caused by PHT pretreatment versus VEH-treatment alone. * Indicates p<0.0001 by Fisher’s exact test. (B) PHT+PTZ treatment groups were divided into immediate (0 minutes post-seizure onset) and delayed (30 minutes post-seizure onset) intervention groups for subsequent pharmacology studies. The majority of PHT+PTZ-treated mice that presented with CSA and were candidates for delayed intervention studies survived for at least 30 min after seizure onset (64/118; 54%). Of the PHT+PTZ mice that were enrolled for delayed intervention group, 50% of mice died within 10 minutes of CSA onset and were thus unable to be rescued with any intervention.

### Continuous Seizure Activity Is Sensitive to Administration of Both Lorazepam and Valproic Acid

We sought to characterize the sensitivity of CSA to both immediate (0 min) and delayed (30 min) interventions using standard-of-care ASDs: LZP, a frontline rescue BDZ for both seizure clusters and SE;^12; 40^ VPA, often utilized as a second line treatment for BDZ-resistant SE;^1^ and CBZ, which can aggravate seizure clusters in humans^16; 41; 42^ and rodent models.^24^ Pharmacological intervention within 1 minute of CSA onset reduced total time spent in CSA (F_(3,61)_=195.7, p<0.0001). However, only LZP and VPA significantly reduced the time spent in CSA (p<0.0001; Figure 3B). Conversely, immediate administration of escalating doses of CBZ did not impact CSA duration. Immediate intervention with high doses of both LZP and VPA significantly improved survival from the CSA insult (Table 1). Finally, immediate administration of all rescue therapies significantly attenuated CSA-induced 24-hour body weight loss versus vehicle-treated mice (F_(3,27)_=3.750, p=0.0225), however only surviving mice were weighed at this point and there was significant mortality in VEH-treated mice (Table 1). Therefore, immediate intervention with both LZP and VPA markedly reduced CSA duration, improved survival from the insult, and reduced CSA-induced body weight loss, whereas immediate intervention with CBZ did not improve any outcome measures aside from improving 24-hour body weight.

**Figure 3.**
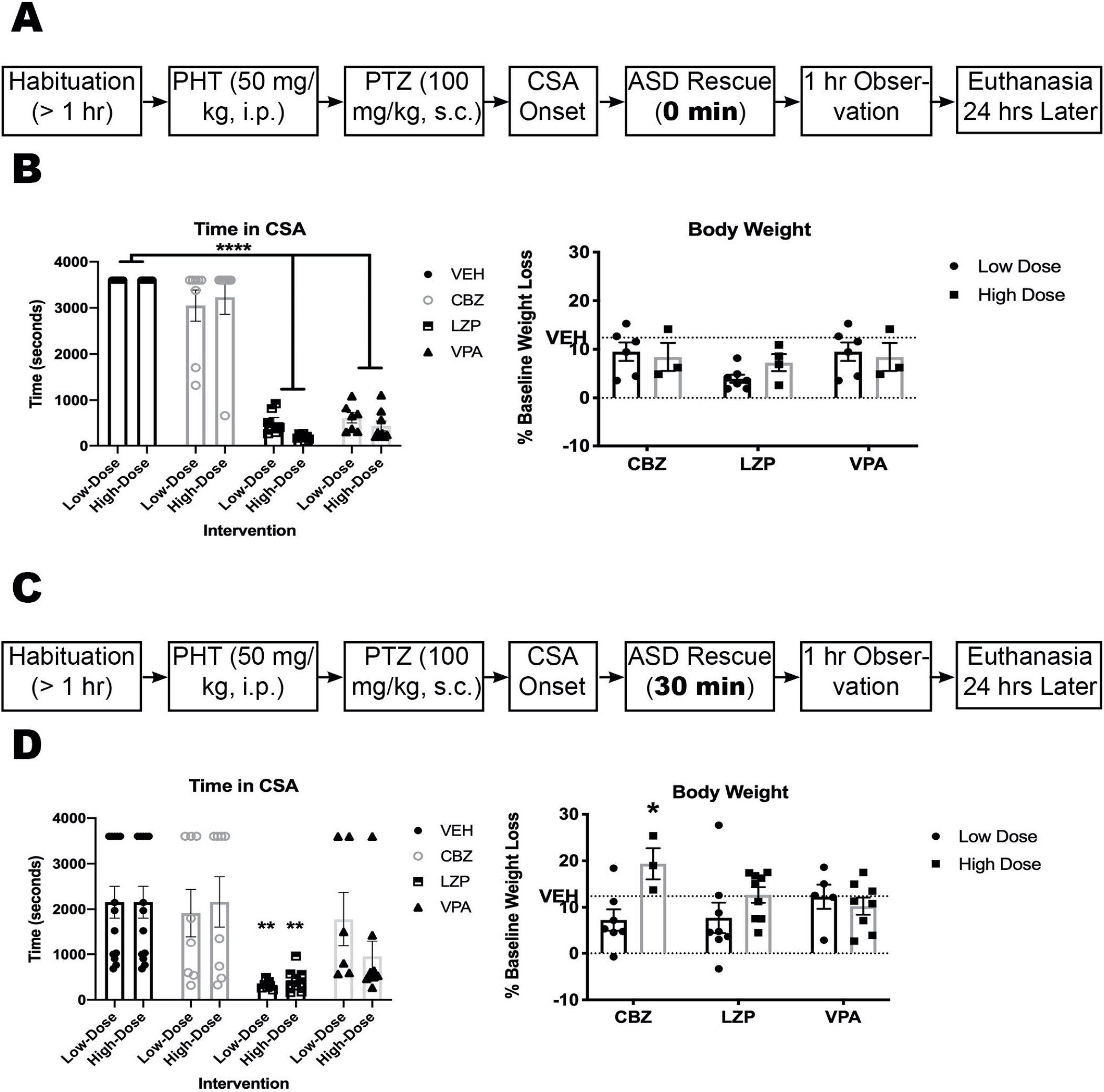
Administration of the antiseizure drugs (ASDs), lorazepam (LZP) and valproic acid (VPA), but not carbamazepine (CBZ), reduce the duration of phenytoin (PHT) + pentylenetetrazol (PTZ)-induced continuous seizure activity (CSA) when administered immediately (0 minutes; A-B) after CSA onset. LZP also reduced CSA duration when administered 30 minutes after CSA onset (C-D) relative to mice treated with vehicle. (A) The timeline of the immediate intervention paradigm to define the pharmacological efficacy of ASDs in this model of seizure clusters. (B) There was a significant effect of treatment on CSA duration in mice treated with both doses of LZP [2, 4 mg/kg] and VPA [150, 300 mg/kg] versus vehicle (F_(3,61)_=195.7, p<0.0001). Administration of CBZ immediately after seizure onset did not significantly reduce CSA duration at either a low or high dose. Data is shown as mean percent body weight loss [+/- SEM] in 24 hours. **** Indicates significantly different from VEH-treated mice, p<0.0001. (C) The timeline of the delayed intervention paradigm to define the pharmacological efficacy of antiseizure drugs in this model of seizure clusters. (D) There was a significant effect of ASD treatment on CSA duration only in mice treated with low- and high-dose LZP versus vehicle (F_(3,66)_=7.246, p=0.0003). Administration of VPA and CBZ 30-minutes after seizure onset did not significantly reduce CSA duration at either a low or high dose. Only LZP effectively reduced seizure duration when administered 30 minutes after CSA onset; ** indicates p<0.01. Data is shown as mean percent body weight loss [+/- SEM] in 24 hours. Further, 40 mg/kg CBZ significantly increased the CSA-induced body weight loss relative to VEH-treated mice; * indicates p=0.037.

Because SE in humans and preclinical models can become refractory to BDZs, we sought to establish whether CSA was also refractory to any intervention administered 30 minutes post-CSA (Figure 3C). Delayed intervention significantly reduced the total time spent in CSA (F_(3,66)_ = 7.246, p=0.0003). However, only LZP reduced the time in CSA in a dose-related fashion (p<0.01; Figure 3D). CBZ and VPA were without significant effect on CSA duration at any dose tested. Delayed intervention with 4 mg/kg LZP also significantly improved survival from CSA (Table 1). There was a main effect of drug dose on 24-hour body weight loss (Figure 3D; F(1,34)=5.20, p=0.029); CBZ (40 mg/kg) was associated with significant worsening of CSA-induced body weight loss versus VEH-treated mice (Figure 3D; p=0.0374) Thus, PHT+PTZ-induced CSA remained sensitive to LZP up to 30 minutes post-onset and administration of this agent alone markedly improved acute CSA-induced behavioral outcomes. Such efficacy of LZP starkly contrasts with traditional SE models that develop BDZ resistance within 30 minutes of seizure onset.

### PHT+PTZ-Induced Electrographic Seizure Activity is Distinct from KA- and s.c.PTZ-Induced Electrographic Seizure Activity

We recorded paired vEEG activity from the PHT+PTZ-treated mice in order to characterize the electrographic seizure profile of a mouse in CSA to more clearly define whether these events were electrographically like those of KA-induced SE or s.c.PTZ-induced myoclonic seizures (Figure 4A). Paired vEEG from mice treated with VEH+PTZ (s.c.PTZ), KA (SE), and PHT+PTZ (CSA) demonstrated qualitatively distinct waveforms in the active seizure period (Figure 4B). Mice treated with VEH+PTZ showed acute electrographic patterns consistent with the behavioral hindlimb extension seizures prototypical of high dose s.c.PTZ (Figure 4C). Mice in KA-induced SE showed a rhythmic pattern of electrographic bursting (Figure 4D) consistent with rodent SE.^23; 34^ Mice in CSA experienced continual electrographic activity with consistent dyssynchronous spiking throughout the recording, distinct from the patterns of KA-induced SE and s.c.PTZ alone (Figure 4E). All groups showed a 10-fold increase in EEG power during the active seizure when compared to their respective baseline, but the small group sizes of this qualitative study did not support statistical comparisons (Figure 4F-H). Nonetheless, there were clear behavioral and electrographic distinctions between s.c.PTZ, KA-SE, and CSA.

**Figure 4.**
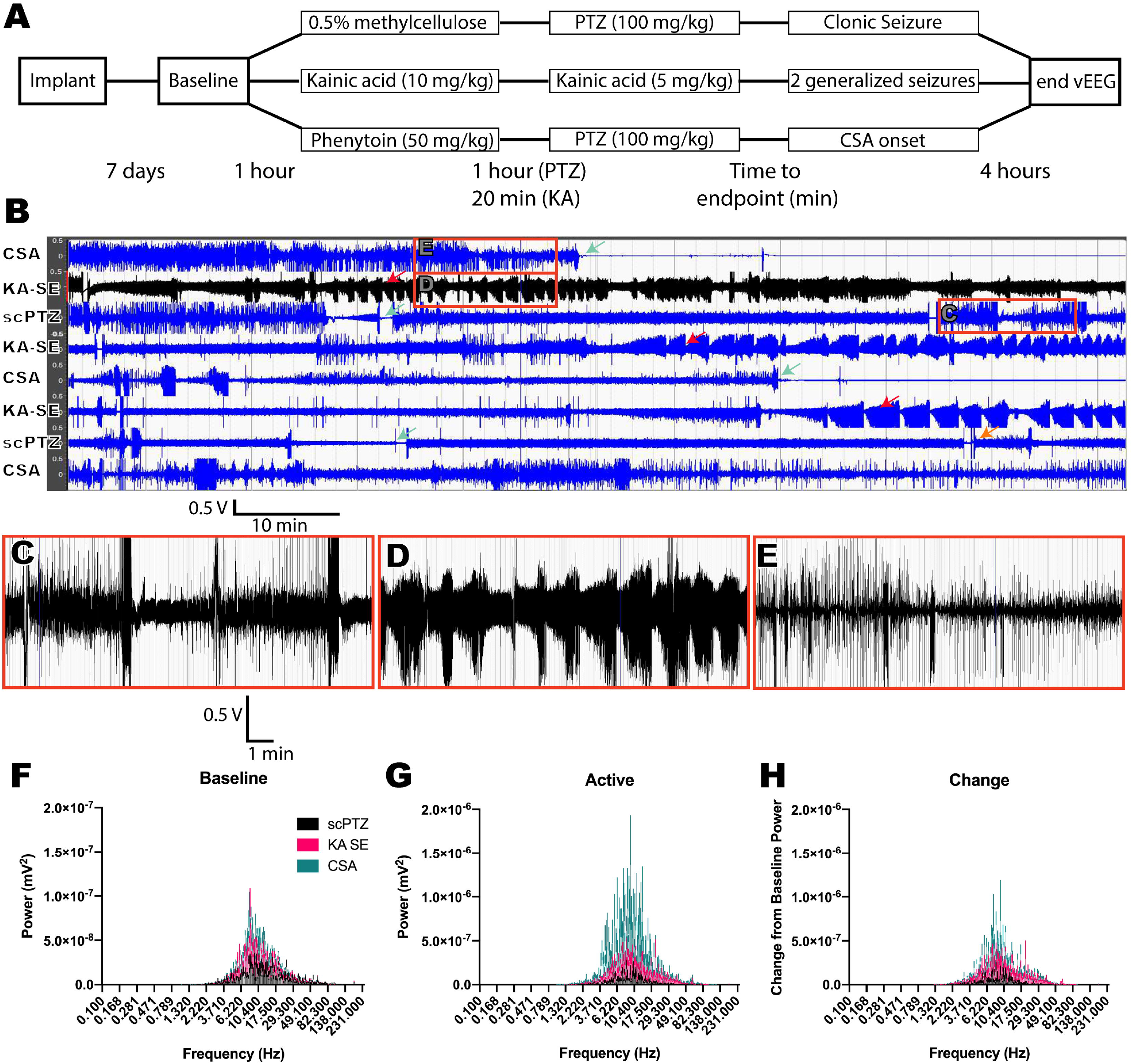
A novel mouse model of continuous seizure activity was produced through the combined administration of PHT (50 mg/kg, i.p.) 1 hour prior to PTZ (s.c. 100 mg/kg) administration. PHT+PTZ produces electrographic patterns visually distinct from those produced during KA-induced status epilepticus (SE). (A) This timeline displays the three distinct experimental flows of the clonic seizure model with VEH+PTZ, the SE model with KA, and the CSA model with PHT+PTZ. (B) 100 minutes of active EEG with each of the three seizure types. Green arrows indicate time of death during a seizure. Red arrows indicate mice in SE following repeated KA administration. Orange arrow indicates a mouse that replaced a deceased animal and that was pretreated with 0.5% methylcellulose vehicle prior to scPTZ administration but never presented with a clonic spasm during the subsequent observation period. Scale bar for 100 minutes of recording. (C-E) 10 minutes of active EEG corresponding to an s.c.PTZ induced clonic hindlimb extension seizure (C), KA induced generalized seizures (D), and PHT+PTZ induced CSA (E). (F) Average baseline power at all frequencies for each testing group (n = 2 s.c.PTZ; n =3 KA SE; n = 3 CSA mice). (G) Average power for the active seizing time measured for a 10-minute period following the onset of a clonic seizure induced by s.c.PTZ, a racine stage 4 or 5 induced by KA, or CSA induced by PHT+PTZ. The y-axis scale shows over a ten-fold increase in power during the active seizing period. (H) The change in EEG power from the baseline recording to active seizing recording periods.

### Continuous Seizure Activity Does Not Induce Neurodegeneration

We quantified hippocampal neurodegeneration with FJ-C staining of tissues collected 24 hours after visually confirmed behavioral CSA onset (n=16) versus tissues obtained from visually confirmed KA-induced SE (n=8 surviving/10 treated mice). KA-SE mice demonstrated extensive neurodegeneration in CA1, CA3, and DG of dorsal hippocampus, as well as neurodegeneration in hypothalamic nucleus (Figure 5A-E). No neurodegeneration was detected in any mice with behavioral CSA (Figure 5F-J). No mice that entered CSA showed any neurodegeneration while every mouse treated with KA had significant neurodegeneration detectable 24 hours after seizure onset (Supplemental Table 1).

**Figure 5.**
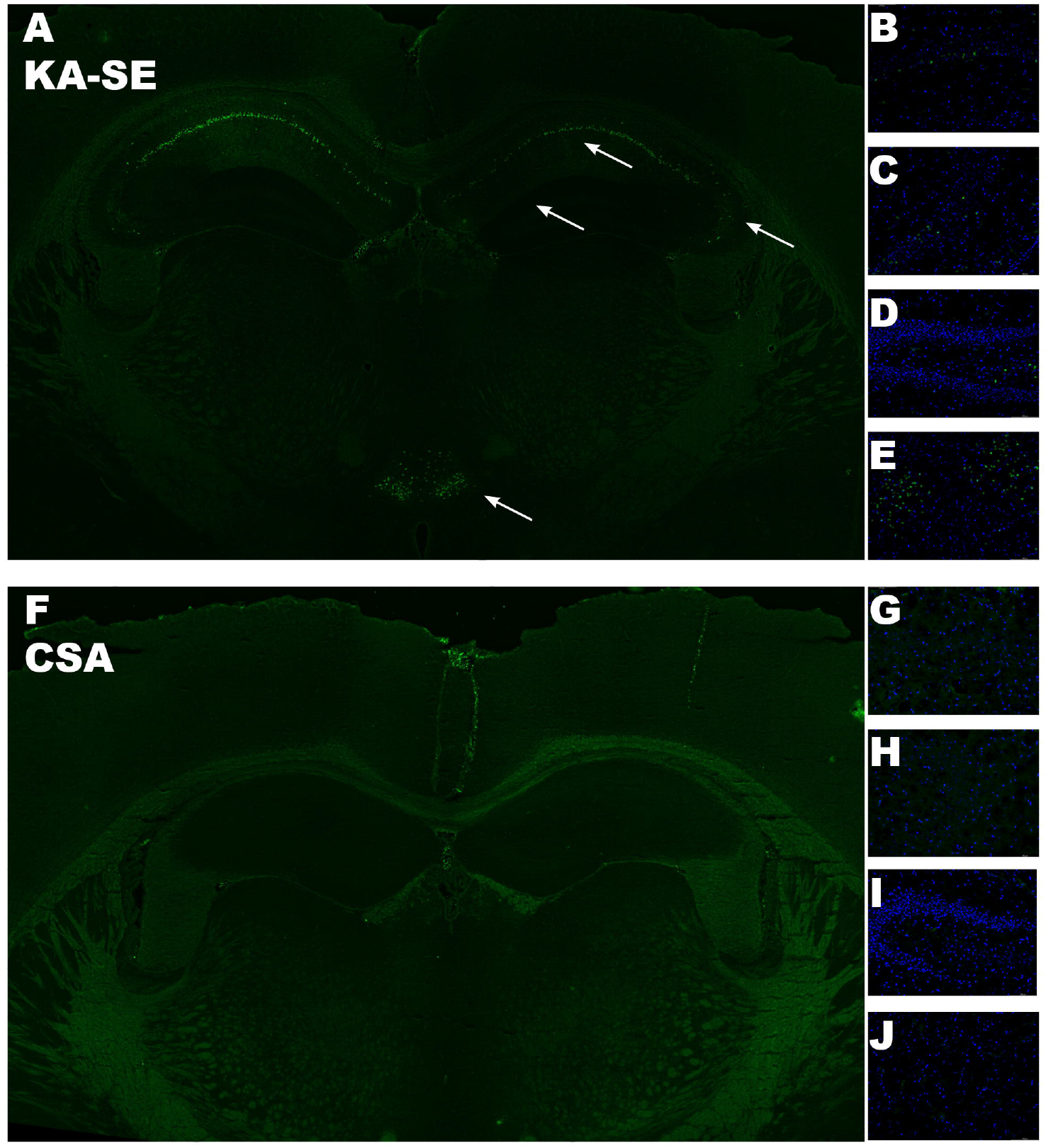
Fluorojade-C (FJ-C) staining of coronal sections from KA-SE and CSA mice collected 24-hours after insult demonstrates that CSA does not induce any neurodegeneration in any limbic structure examined. (A) FJ-C stained whole brain slice showing neurodegeneration in the hippocampus and hypothalamic nucleus of a CF-1 mouse that underwent KA induced SE. Arrows highlight areas of FJ-C positive cells, indicative of dead and dying neurons. (B-E) There is widespread and distinct neurodegeneration in B) CA1, C) CA3, D) dentate gyrus, and E) hypothalamic nucleus of a mouse following KA-induced SE. (F) FJ-C stained whole brain slice showing no neurodegeneration in the hippocampus and hypothalamic nucleus of a CF-1 mouse that underwent PHT+PTZ-induced CSA. (G-J) There are no dead and dying neurons present in the G) CA1, H) CA3, I) dentate gyrus, or J) hypothalamic nucleus following PHT+PTZ-induced CSA.

## Discussion

SE is a distinct seizure emergency characterized by prolonged seizures lasting over 30 minutes with no recovery. BDZs are the standard-of-care, but SE can quickly become refractory to BDZs if untreated, increasing the risk of morbidity and mortality. We presently demonstrate that mice pretreated with high dose PHT (50 mg/kg) prior to s.c.PTZ (100 mg/kg) exhibit a seizure phenotype distinct from the hindlimb extension seizures characteristic of s.c.PTZ.^27^ These seizures are unequivocally distinct from KA-induced SE at the behavioral, pharmacological, electrographical, and neuropathological levels. CSA is thus a unique model of seizure emergencies.

The precise mechanisms that contribute to the presently described onset of CSA following PHT+PTZ remain to be elucidated but provide a unique opportunity to further interrogate the processes underlying seizure termination. PHT, an ASD that blocks sodium channels, does not prevent minimal clonic seizures induced by s.c.PTZ, even at PHT doses above 100 mg/kg, but PHT effectively prevents maximal hindlimb tonic extension seizures resulting from suprathreshold doses of s.c.PTZ.^27^ PHT likely blocks seizure spread but does not increase seizure threshold, in contrast to other ASDs.^27^ The NINDS ETSP has even demonstrated that PHT (>41 mg/kg) can actually decrease seizure threshold in the mouse i.v.PTZ test (https://panache.ninds.nih.gov). Consistent with earlier reports,^29^ we herein demonstrate that pretreatment with high dose PHT (50 mg/kg) shifts the phenotypic seizure of high dose s.c.PTZ-induced seizures from hindlimb tonic extension to sustained CSA. PHT+PTZ promotes CSA in over 73% of mice (of n=244), a phenomenon that is best described as sustained clonic seizures. Conversely, only 14% of VEH+PTZ-treated mice (of n=94) demonstrated CSA; the majority exhibited hindlimb extension seizures typical of suprathreshold doses of s.c.PTZ. The fact that PHT prevents seizure spread and blocks tonic hindlimb extension but reduces seizure threshold at high (i.e., proconvulsant) doses likely contributed to the CSA phenomenon when PHT was administered in combination with s.c.PTZ. Seizures in patients and preclinical models will generally self-terminate within 1-2 minutes and seizure duration is relatively short.^43^ Until now, this statement generally held true for all events except SE, which likely explains the rationale for a prior study wherein combined administration of PHT+PTZ was used to explore the efficacy of midazolam for SE.^29^ CSA in mice is distinguished from KA-induced SE based on electrographic, behavioral, and pharmacological features, demonstrating that our model is not only distinct from acute seizures and SE, but that it is also a novel platform on which to elucidate the mechanisms contributing to seizure termination.

Seizure emergencies are clinically managed with BDZs.^44^ The activity of LZP, a first line therapy for both seizure clusters and SE,^9; 12; 40^ administered with a 30-minute delay demonstrates that CSA is not refractory to BDZs.^29^ CSA is highly sensitive to LZP at otherwise refractory time points (30 minutes – our study; or longer^29^). Rodent models of SE are refractory to BDZs consistent with clinical SE.^1; 45^ Further, VPA, a common second line therapy for SE and seizure clusters,^46–48^ was as effective as LZP on both reducing CSA duration and improving 24-hour survival following immediate administration (Figure 3B) but lost efficacy by 30 minutes (Figure 3D). Efficacy of these two pharmacologically distinct, clinically effective ASDs supports the face validity of this model as a new tool to interrogate the mechanisms underlying the onset and termination of seizure emergencies.

CSA could also be worsened by ASD intervention. Specifically, administration of CBZ 30 minutes after onset did not reduce CSA duration and actually reduced survival and increased 24-hour body weight loss. These findings align with clinical reports wherein CBZ and other sodium channel blockers can aggravate seizure clusters in patients^16; 41; 42^□ as well as in a rat model of temporal lobe epilepsy with documented spontaneous seizure clusters^24^□ Thus, CBZ confirmed that CSA was bidirectionally sensitive to ASDs, consistent with seizure clusters. Taken together the findings that LZP (our study) and other BDZs^29^□ and that CBZ worsens outcomes, in our current investigation clearly indicates that the observed phenomenon associated with PHT and s.c.PTZ coadministration is not representative of SE, but rather another seizure emergency; i.e., non-SE CSA.

Systemic administration of KA to rodents is an established method to induce electrographic seizures that progress to rhythmic recurrent electrographic spiking and convulsive SE; these events are detectable by cortical EEG. While the systemic low-dose KA-induced SE model has been previously reported in inbred C57Bl/6 mice,^34^ we now established this model in outbred CF-1 mice and find it to be a reliable model of acute SE with clear and consistent, rhythmic spiking. However, mice that underwent CSA did not demonstrate such rhythmic firing patterns (Figure 4). KA-SE also leads to severe and dispersed neurodegeneration throughout numerous limbic structures, as assessed by FJ-C staining for degenerating neurons, in all mice 24-hours after insult; none of the CSA mice demonstrated any evidence of neurodegeneration in any examined brain region 24-hours later (Figure 5). Altogether, the electrographic seizure patterns, changes in EEG power between experimental groups, and the clear lack of neurodegeneration 24-hours after insult confirm that this CSA model induces a sustained seizure insult that is altogether pathologically distinct from SE.

The CSA model provides a new platform for the study of mechanisms underlying seizure emergencies, as well as their differences in ASD sensitivity. The majority of mice that entered CSA survived for at least 30 minutes post-insult, allowing for interventions to be administered at an investigator-defined time point in a manner similar to preclinical SE models.^49^ Whether BDZs are the only and best-suited interventions for all seizure emergencies has not yet been rigorously evaluated in a preclinical model. There may be better-tolerated or more effective interventions that are specific to SE versus seizure clusters. Further, this study establishes the CSA model as a platform on which to identify other potential treatments, even with delayed administration, to rescue seizure emergencies. Rescue therapies for seizure emergencies are not often readily available in an outpatient setting; the timing of drug administration may be long after seizure onset.^50^ In this regard, preclinical models with sustained seizures that remain sensitive to protracted interventions could be highly informative to drug discovery and differentiation. Whether this CSA model will ultimately prove useful in the identification of novel interventional agents for the management of seizure emergencies clearly remains to be established.

## Acknowledgements

We confirm that we have read the Journal’s position on issues involved in ethical publication and affirm that this report is consistent with those guidelines. This work was supported by the University of Washington Department of Pharmacy and the NCATS/Institute for Translational Health Sciences (NCATS KL2 TR002317 to MBH). The authors are grateful for the curated list of clinical seizure studies provided by Saifuddin Kharawala and UCB Pharma as AES Abstract 3.331 (December 2019; Baltimore, MD). The authors acknowledge the technical assistance of Zachery Koneval and graphical design assistance of Matthew Haliski.

## References

1. Trinka E, Shorvon S. A decade of progress in status epilepticus 2007-2017: Proceedings of the 6th London-Innsbruck Colloquium on Status Epilepticus and Acute Seizures. Epilepsia 2018;59 Suppl 2:67–69.

2. Hauser WA. Status epilepticus: epidemiologic considerations. Neurology 1990;40:9–13.

3. Neligan A, Noyce AJ, Gosavi TD, et al. Change in Mortality of Generalized Convulsive Status Epilepticus in High-Income Countries Over Time: A Systematic Review and Meta-analysis. JAMA Neurol 2019.

4. Hauser WA, Rich SS, Annegers JF, et al. Seizure recurrence after a 1st unprovoked seizure: an extended follow-up. Neurology 1990;40:1163–1170.

5. Theodore WH, Porter RJ, Albert P, et al. The secondarily generalized tonic-clonic seizure: a videotape analysis. Neurology 1994;44:1403–1407.

6. Jenssen S, Gracely EJ, Sperling MR. How long do most seizures last? A systematic comparison of seizures recorded in the epilepsy monitoring unit. Epilepsia 2006;47:1499–1503.

7. Rose AB, McCabe PH, Gilliam FG, et al. Occurrence of seizure clusters and status epilepticus during inpatient video-EEG monitoring. Neurology 2003;60:975–978.

8. Wallace SJ. Nasal benzodiazepines for management of acute childhood seizures? Lancet 1997;349:222.

9. Silbergleit R, Durkalski V, Lowenstein D, et al. Intramuscular versus intravenous therapy for prehospital status epilepticus. N Engl J Med 2012;366:591–600.

10. Nunley S, Glynn P, Rust S, et al. A hospital-based study on caregiver preferences on acute seizure rescue medications in pediatric patients with epilepsy: Intranasal midazolam versus rectal diazepam. Epilepsy Behav 2019;92:53–56.

11. Owusu KA, Dhakar MB, Bautista C, et al. Comparison of intranasal midazolam versus intravenous lorazepam for seizure termination and prevention of seizure clusters in the adult epilepsy monitoring unit. Epilepsy Behav 2019;98:161–167.

12. Mitchell WG. Status epilepticus and acute repetitive seizures in children, adolescents, and young adults: etiology, outcome, and treatment. Epilepsia 1996;37 Suppl 1:S74–80.

13. Mayer SA, Claassen J, Lokin J, et al. Refractory status epilepticus: frequency, risk factors, and impact on outcome. Arch Neurol 2002;59:205–210.

14. Kutlu NO, Yakinci C, Dogrul M, et al. Intranasal midazolam for prolonged convulsive seizures. Brain Dev 2000;22:359–361.

15. Cereghino JJ. Identification and treatment of acute repetitive seizures in children and adults. Curr Treat Options Neurol 2007;9:249–255.

16. Haut SR, Shinnar S, Moshe SL. Seizure clustering: risks and outcomes. Epilepsia 2005;46:146–149.

17. Haut SR, Shinnar S, Moshe SL, et al. The association between seizure clustering and convulsive status epilepticus in patients with intractable complex partial seizures. Epilepsia 1999;40:1832–1834.

18. Barker-Haliski M, Harte-Hargrove LC, Ravizza T, et al. A companion to the preclinical common data elements for pharmacologic studies in animal models of seizures and epilepsy. A Report of the TASK3 Pharmacology Working Group of the ILAE/AES Joint Translational Task Force. Epilepsia Open 2018;3:53–68.

19. Loscher W. Fit for purpose application of currently existing animal models in the discovery of novel epilepsy therapies. Epilepsy Res 2016;126:157–184.

20. Sperk G. Kainic acid seizures in the rat. Prog Neurobiol 1994;42:1–32.

21. Rogawski MA, Bazil CW. New molecular targets for antiepileptic drugs: alpha(2)delta, SV2A, and K(v)7/KCNQ/M potassium channels. Curr Neurol Neurosci Rep 2008;8:345–352.

22. McKhann GM, 2nd, Wenzel HJ, Robbins CA, et al. Mouse strain differences in kainic acid sensitivity, seizure behavior, mortality, and hippocampal pathology. Neuroscience 2003;122:551–561.

23. Saporito MS, Gruner JA, DiCamillo A, et al. Intravenously Administered Ganaxolone Blocks Diazepam-Resistant Lithium-Pilocarpine-Induced Status Epilepticus in Rats: Comparison with Allopregnanolone. J Pharmacol Exp Ther 2019;368:326–337.

24. Grabenstatter HL, Dudek FE. Effect of carbamazepine on spontaneous recurrent seizures recorded from the dentate gyrus in rats with kainate-induced epilepsy. Epilepsia 2019;60:636–647.

25. Williams PA, White AM, Clark S, et al. Development of spontaneous recurrent seizures after kainate-induced status epilepticus. J Neurosci 2009;29:2103–2112.

26. Mizuno S, Koneval Z, Zierath DK, et al. Diurnal Burden of Spontaneous Seizures in Early Epileptogenesis in the Post-Kainic Acid Rat Model of Epilepsy *Epilepsia Open* 2021;In Press.

27. Piredda SG, Woodhead JH, Swinyard EA. Effect of stimulus intensity on the profile of anticonvulsant activity of phenytoin, ethosuximide and valproate. J. Pharmacol. Exp. Ther. 1985;232:741–745.

28. Loscher W. Preclinical assessment of proconvulsant drug activity and its relevance for predicting adverse events in humans. Eur J Pharmacol 2009;610:1–11.

29. Raines A, Henderson TR, Swinyard EA, et al. Comparison of midazolam and diazepam by the intramuscular route for the control of seizures in a mouse model of status epilepticus. Epilepsia 1990;31:313–317.

30. Kilkenny C, Browne W, Cuthill IC, et al. Animal research: reporting in vivo experiments--the ARRIVE guidelines. J Cereb Blood Flow Metab 2011;31:991–993.

31. White HS, Woodhead JH, Wilcox KS, et al. Discovery and preclinical development of antiepileptic drugs. In Levy RH, Mattson RH, Meldrum BS, et al. (Eds) Antiepileptic Drugs, Fifth Edition, Lippincott Williams & Wilkins: Philadelphia; 2002:36–48.

32. Bialer M, Twyman RE, White HS. Correlation analysis between anticonvulsant ED50 values of antiepileptic drugs in mice and rats and their therapeutic doses and plasma levels. Epilepsy Behav 2004;5:866–872.

33. Barker-Haliski ML, Johnson K, Billingsley P, et al. Validation of a Preclinical Drug Screening Platform for Pharmacoresistant Epilepsy. Neurochem Res 2017;42:1904–1918.

34. Umpierre AD, Bennett IV, Nebeker LD, et al. Repeated low-dose kainate administration in C57BL/6J mice produces temporal lobe epilepsy pathology but infrequent spontaneous seizures. Exp Neurol 2016;279:116–126.

35. Stewart KA, Wilcox KS, Fujinami RS, et al. Development of postinfection epilepsy after Theiler’s virus infection of C57BL/6 mice. J Neuropathol Exp Neurol 2010;69:1210–1219.

36. Schmued LC, Stowers CC, Scallet AC, et al. Fluoro-Jade C results in ultra high resolution and contrast labeling of degenerating neurons. Brain Res 2005;1035:24–31.

37. Wanisch K, Kovac S, Schorge S. Tackling obstacles for gene therapy targeting neurons: disrupting perineural nets with hyaluronidase improves transduction. PLoS One 2013;8:e53269.

38. Finney DJ. Probit analysis. A statistical treatment of the sigmoid response curve. University Press: Cambridge; 1952.

39. Litchfield JR, Jr., Wilcoxon R. A simplified method of evaluating dose-effect experiments. J. Pharmacol. 1949;96:99–113.

40. Treiman DM, Meyers PD, Walton NY, et al. A comparison of four treatments for generalized convulsive status epilepticus. Veterans Affairs Status Epilepticus Cooperative Study Group. N Engl J Med 1998;339:792–798.

41. Haut SR. Seizure clustering. Epilepsy Behav 2006;8:50–55.

42. Bauer J, Ghane Y, Flugel D, et al. [Etiology, follow-up and therapy of seizure clusters in temporal lobe epilepsy and catamenial epileptic seizures]. Schweiz Arch Neurol Psychiatr (1985) 1992;143:117–134.

43. Loscher W, Kohling R. Functional, metabolic, and synaptic changes after seizures as potential targets for antiepileptic therapy. Epilepsy Behav 2010;19:105–113.

44. Jafarpour S, Hirsch LJ, Gainza-Lein M, et al. Seizure cluster: Definition, prevalence, consequences, and management. Seizure 2019;68:9–15.

45. Naylor DE, Liu H, Niquet J, et al. Rapid surface accumulation of NMDA receptors increases glutamatergic excitation during status epilepticus. Neurobiol Dis 2013;54:225–238.

46. Gilad R, Izkovitz N, Dabby R, et al. Treatment of status epilepticus and acute repetitive seizures with i.v. valproic acid vs phenytoin. Acta Neurol Scand 2008;118:296–300.

47. Brigo F, Storti M, Del Felice A, et al. IV Valproate in generalized convulsive status epilepticus: a systematic review. Eur J Neurol 2012;19:1180–1191.

48. Brophy GM, Bell R, Claassen J, et al. Guidelines for the evaluation and management of status epilepticus. Neurocrit Care 2012;17:3–23.

49. Loscher W. Strategies for antiepileptogenesis: Antiepileptic drugs versus novel approaches evaluated in post-status epilepticus models of temporal lobe epilepsy. In Noebels JL, Avoli M, Rogawski MA, et al. (Eds) Jasper’s Basic Mechanisms of the Epilepsies: Bethesda (MD); 2012.

50. Penovich PE, Buelow J, Steinberg K, et al. Burden of Seizure Clusters on Patients With Epilepsy and Caregivers: Survey of Patient, Caregiver, and Clinician Perspectives. Neurologist 2017;22:207–214.

