## Supplementary material for "Continuous Seizure Emergency Evoked in Mice with Pharmacological, Electrographic, and Pathological Features Distinct from Status Epilepticus": Table 1

Table 1. 24-hour Survival

|  | Vehicle | CBZ  20 mg/kg | CBZ  40 mg/kg | LZP  2 mg/kg | LZP  4 mg/kg | VPA  150 mg/kg | VPA  300 mg/kg |
| --- | --- | --- | --- | --- | --- | --- | --- |
| Immediate Intervention | 5/11 | 6/8 | 3/8 | 7/8 | 8/8 | 6/8 | 9/9 |
| Delayed Intervention | 12/13 | 7/8 | 3/8 | 8/8 | 9/9 | 6/8 | 8/9 |
