## Supplemental Table 1 for "Continuous Seizure Emergency Evoked in Mice with Pharmacological, Electrographic, and Pathological Features Distinct from Status Epilepticus"

|  | Presence of FJ-C Positive cells in CA1 | Presence of FJ-C positive cells in CA3 | Presence of FJ-C positive cells in the Dentate Gyrus | Presence of FJ-C positive cells in the Hypothalamic Nucleus |
| --- | --- | --- | --- | --- |
| PHT+PTZ induced CSA rescued with Vehicle | 0/16 | 0/16 | 0/16 | 0/16 |
| Kainic Acid induced SE | 4/8 *, p=0.0066 | 5/8 *, p = 0.0013 | 6/8 *, p=0.0002 | 7/8 *, p<0.0001 |

*significantly different from CSA mice by Fisher’s exact two-sided test
